# Highly-parallel microfluidics-based force spectroscopy on single cytoskeletal motors

**DOI:** 10.1101/2020.08.11.245910

**Authors:** Marta Urbanska, Annemarie Lüdecke, Wim J. Walter, Antoine M. van Oijen, Karl E. Duderstadt, Stefan Diez

**Author notes:** correspondence and requests for materials should be addressed to K.E.D. and S.D. Max Planck Institute for the Science of Light & Max-Planck-Zentrum für Physik und Medizin, 91058 Erlangen, Germany (Marta Urbanska). NanoTemper Technologies GmbH, 81369 München, Germany (Annemarie Lüdecke). Health and Medical University Potsdam, 14471 Potsdam, Germany (Wim J. Walter).

## Abstract

Cytoskeletal motors transform chemical energy into mechanical work to drive essential cellular functions. Optical trapping experiments have provided crucial insights into the operation of these molecular machines under load. However, the throughput of such force spectroscopy experiments is typically limited to one measurement at a time. Here, we introduce a highly-parallel, microfluidics-based method that allows for rapid collection of force-dependent motility parameters of cytoskeletal motors with two orders of magnitude improvement in throughput compared to currently available methods. We apply tunable hydrodynamic forces to stepping kinesin-1 motors via DNA-tethered beads and utilize a large field of view to simultaneously track the velocities, run lengths and interaction times of hundreds of individual kinesin-1 molecules under varying resisting and assisting loads. Importantly, the 16-μm long DNA tethers between the motors and the beads significantly reduces the vertical component of the applied force pulling the motors away from the microtubule. Our approach is readily applicable to other molecular systems and constitutes a new methodology for parallelized single-molecule force studies on cytoskeletal motors.

## 1. Introduction

The application and detection of forces using single-molecule manipulation methods has provided major advances in our understanding of the operating principles of mechanoenzymes.^[1–4]^ Optical and magnetic tweezers as well as atomic force microscopy are now being routinely used to study protein folding pathways, receptor-ligand interactions, DNA mechanics and the activity of molecular motors. While all of these experimental approaches offer excellent spatiotemporal resolution and force accuracy—with different force spectra and displacement ranges covered—none of them provides high experimental throughput as conventionally only one molecule is studied at a time. This limitation constitutes one of the major bottlenecks in current single-molecule force measurements,^[4,5]^ where the derivation of statistically significant results from stochastic single-molecule footprints is desired in reasonable time frames. Consequently, continuous efforts are being made to surpass this limitation in the field of optical^[6–8]^ and magnetic trapping^[9–11]^ as well as in atomic force microscopy—regarding both instrumental automation^[12]^ as well as sample preparation.^[13]^ Alongside, a number of novel solutions for multiplexed force manipulation, such as centrifuge force microscopy^[14,15]^ and acoustic force spectroscopy,^[16]^ are being introduced. So far, the use of these novel methods has been demonstrated for the studies of DNA mechanics, DNA-protein binding and protein-protein binding but not for cytoskeletal motor proteins. While their use to study DNA motors is conceivable, they may require modifications to become applicable for studies on cytoskeletal motors because of the vertical character of the applied force.

Cytoskeletal motor proteins generate forces using the energy from adenosine triphosphate (ATP) hydrolysis to move along filamentous tracks and execute cellular functions such as cargo transportation or chromosome separation in mitosis.^[17]^ The discovery of the microtubule-associated motor protein kinesin-1,^[18]^ the major workhorse of long-range anterograde cargo transport, was followed shortly by its manipulations with optical tweezers.^[19–21]^ In the optical tweezer experiments, the influence of force on the motility parameters that characterize the movement of individual motors, including run length, stepping velocity and interaction time, can be investigated to gain insights into the mechanics of the motor operation. Run length, a hallmark of motor processivity, refers to the distance covered by each kinesin molecule before detaching from a microtubule, stepping velocity describes the pace at which the motor translocates, and interaction time is the time spent on the microtubule between landing and dissociation events. Even though originally several ways of exerting load on stepping kinesins such as glass fiber manipulation,^[22]^ microtubule bending^[23]^ and viscous load^[24]^ were exploited, at present optical tweezers are used almost exclusively for the characterization of cytoskeletal motors under loads. Apart from offering comparatively low throughput, it has been argued that the poor control over force applied in the vertical direction in the optical tweezers setup may bias the measurements performed with this method.^[25,26]^

One so far largely unexploited way to apply calibrated forces onto individual molecules is hydrodynamic flow. In a microfluidic environment, laminar flow can be used to exert Stokes drag on micrometer-sized beads that act as force handles when linked to surface-attached biomolecular mechanosystems. The magnitude of the drag force is determined by the diameter of the beads and the velocity of the flow. The latter can be kept constant over large regions in a microfluidic chamber. The response of the molecular system under investigation can then be deduced by tracking the positions of multiple beads simultaneously using an optical microscope. Hence, the number of constant-force experiments performed at a time is in principle limited only by the size of the imaged area and the surface density of the bead-coupled mechanosystems. Low-throughput experiments using hydrodynamic flow have so far been performed to study single-molecule forces in protein unfolding,^[27]^ to measure rupture forces of streptavidin-biotin bonds,^[28]^ to investigate the confining potential felt by individual membrane-embedded receptors^[29]^ and, in the context of cytoskeletal motors, to measure the adhesion forces of beads covered with multiple kinesin-1 motors to microtubules.^[30]^ Moreover, high-throughput experiments using hydrodynamic flow have been performed to study DNA mechanics and DNA-protein interactions. In particular, highly-parallel measurements to monitor the enzymatic activity of DNA exonucleases,^[31]^ DNA and RNA polymerases,^[32–35]^ or topoisomerases^[36]^ have been demonstrated on flow-stretched DNA, with force control down to 0.1 pN.

Here, we demonstrate the application of hydrodynamic forces to investigate the translocation of cytoskeletal motors under load in a highly parallel manner. In particular, we use paramagnetic beads attached to individual kinesin-1 motors via 16-μm long DNA linkers as force handles and utilized a large field of view microscope to characterize the velocities, run lengths and interaction times of hundreds of motors stepping under a series of in situ calibrated force conditions. Leveraging the large spatially homogenous force field generated by hydrodynamic flow and the use of a specialized telecentric lens capturing a field of view of several millimeters in size, we were able to optically track hundreds of individual molecules in a single experiment, amounting to a total of 2,500 events in eight experiments. Consistent with previous low-throughput measurements with glass fiber^[22]^ and optical tweezers,^[37,38]^ our data shows that the velocity of kinesin-1 motors gradually decreased under increasing load by up to 62% for resisting loads of 3.3 pN. For assisting loads of the same magnitude, the velocity decreased by up to 35%. Due to the molecular geometry of our assay, we were able to directly measure the motility parameters of kinesin-1 in the absence of significant vertical forces (i.e. away from the microtubule surface), which had previously been only accessible by theoretical calculations. Our high-throughput method does not require expensive equipment and can be easily adapted to other biomolecular mechanosystems.

## 2. Results

### 2.1. Molecular assembly of the mechanosystems

To assemble the molecular system for tracking of individual kinesin-1 motors stepping along microtubules under load, we sequentially attached specially designed molecular components to the surface of a flow cell by flowing them through the flow cell using a syringe pump (**Figure 1a** and **Methods**). First, guanosine-5’-[αβ-methyleno] triphosphate (GMPCPP)- stabilized microtubules were immobilized on the surface via anti-β-tubulin antibodies. Next, truncated, SNAP-tagged kinesin-1 motors were covalently coupled to 16-μm long double-stranded DNA (dsDNA) linkers based on lambda phage DNA (λ-DNA) with functionalized ends. The kinesin-DNA complexes were then introduced to the flow cell and attached to the microtubules under flow in the presence of 100 μM adenylyl-imidodiphosphate (AMPPNP). Finally, 1-μm sized superparamagnetic beads coated with anti-digoxigenin antibodies were flowed in and attached to the free ends of the DNA linkers. The AMPPNP kept the kinesin-1 motors at fixed positions on the microtubules until the beginning of the measurement which was initiated by the addition of 10 mM ATP (**Figure S1**). Evaluation of the extreme positions of the beads during flow reversal showed that the length of most tethers corresponds to the full length single λ-DNA (**Figure S2**), indicating prevalence of full-length molecules with single attachment sites.

**Figure 1.**
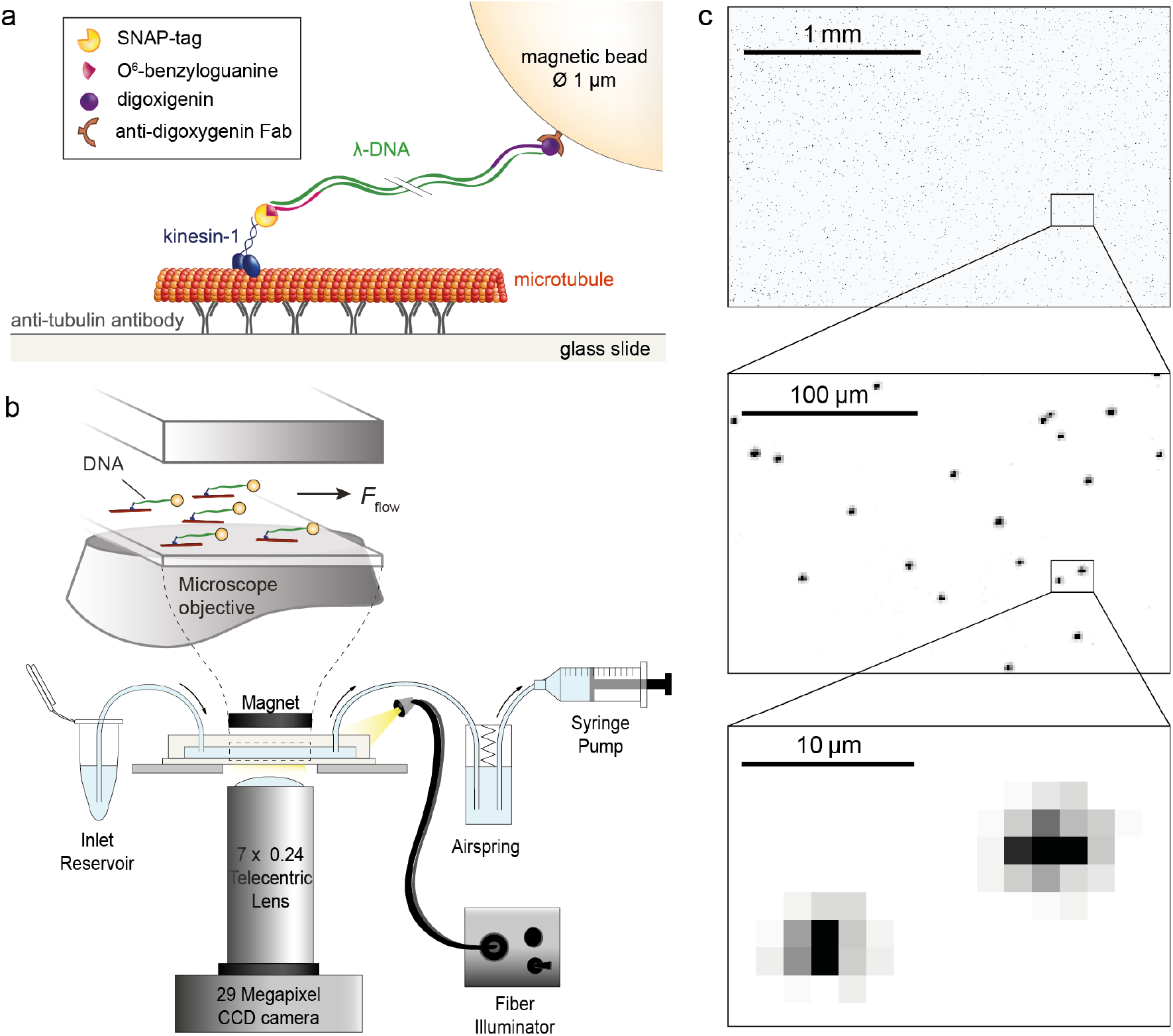
Kinesin-1 microfluidics-based force assay. a) Molecular details of attaching a 1-μm sized paramagnetic bead to an individual kinesin-1 motor via a long, double-stranded DNA linker. b) Schematic overview of the experimental setup. Inset presents a side view of the interior of the flow cell (not to scale). The direction of applied force is indicated by the arrow. c) Dark-field microscopy images of multiple magnetic beads (up to 30,000 beads per field of view) tethered to individual kinesin-1 motors. The top panel represents only 23% of the full field of view.

### 2.2. Microfluidics-based force assay

The heart of the experimental setup constituted a custom-made inverted microscope (**Figure 1b**). A syringe pump, operated in withdrawal mode, was used to apply a hydrodynamic flow throughout the experiment. An air spring was introduced between the flow cell outlet and the pump in order to damp any flow irregularities. To minimize interactions of the tethered beads with the surface, a magnet installed on the top of the flow cell provided a miniscule force of approximately 0.1 pN to lift the paramagnetic beads up. To observe the bead positions, the flow cell was illuminated from the side with high-intensity white light from a fiber illuminator. The light scattered by the beads was collected through a telecentric lens and projected onto a large-format CCD camera. In contrast to conventional lenses, telecentric lenses have a constant, non-angular field of view, i.e. the chief rays are perpendicular to the object plane over the full field of view.^[39]^ This assures the same magnification independent of the distance between the imaged object and the lens, diminishes image distortion towards the image edges and eliminates projection errors that could arise from tilt in the object plane, altogether providing a high accuracy of measured distances over the whole field of view.^[40]^

Due to the large scattering cross-section of the beads it was possible to implement a low-magnification objective to maximize the field of view without substantial loss in accuracy when determining the bead positions. The high quality of the telecentric lens, together with the 29 Megapixel camera sensor, provided for distance-accurate imaging of an 18 mm^2^ large region. Within one field of view it was possible to image up to 30,000 beads (**Figure 1c**) and each of them could be tracked with a precision of 32 nm (see **Methods**).

### 2.3. In situ force calibration

As routinely used in the field of magnetic tweezers, we used the fluctuations of the tethered beads in the direction transverse to the flow for an in situ calibration of the acting forces.^[41]^ By relating the energy of a Hookean spring to the equipartition theorem the following equation is obtained:

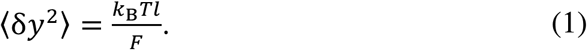

The mean-square displacement of a bead in the transverse direction <*δy*^*2*^>, together with the length of the tether *l*, temperature *T* and Boltzmann constant *k*_*B*_ are sufficient to determine the force *F* pulling on the molecule. To enable precise determination of the tether extension for each molecule, we coupled the force-extension relation for dsDNA^[42]^ with Equation (1) and solved the set of these two equations numerically to obtain both tether extension and force magnitude for each molecule individually (see **Methods** for details, including a correction for motion blurring caused by the finite camera integration time).

The measured magnitude of the fluctuations of individual beads decreased with increasing flow rate (**Figure 2a**). Using the trajectories from all beads which exhibited unidirectional movement after the addition of ATP, we determined a characteristic force for each experiment. **Figure 2b** presents the force distributions for exemplary experiments performed at flow rates of 10, 20, 30 and 40 μl min^-1^. The median forces in the presented experiments were 1.1, 1.7, 2.2 and 2.9 pN, respectively. The broad distribution of estimated forces can be attributed to local flow instabilities, as well as to a potential non-uniformity of bead sizes. The influence of the latter could, in the future, be assessed by bead sizing using convolution and correlation image analysis.^[43]^

**Figure 2.**
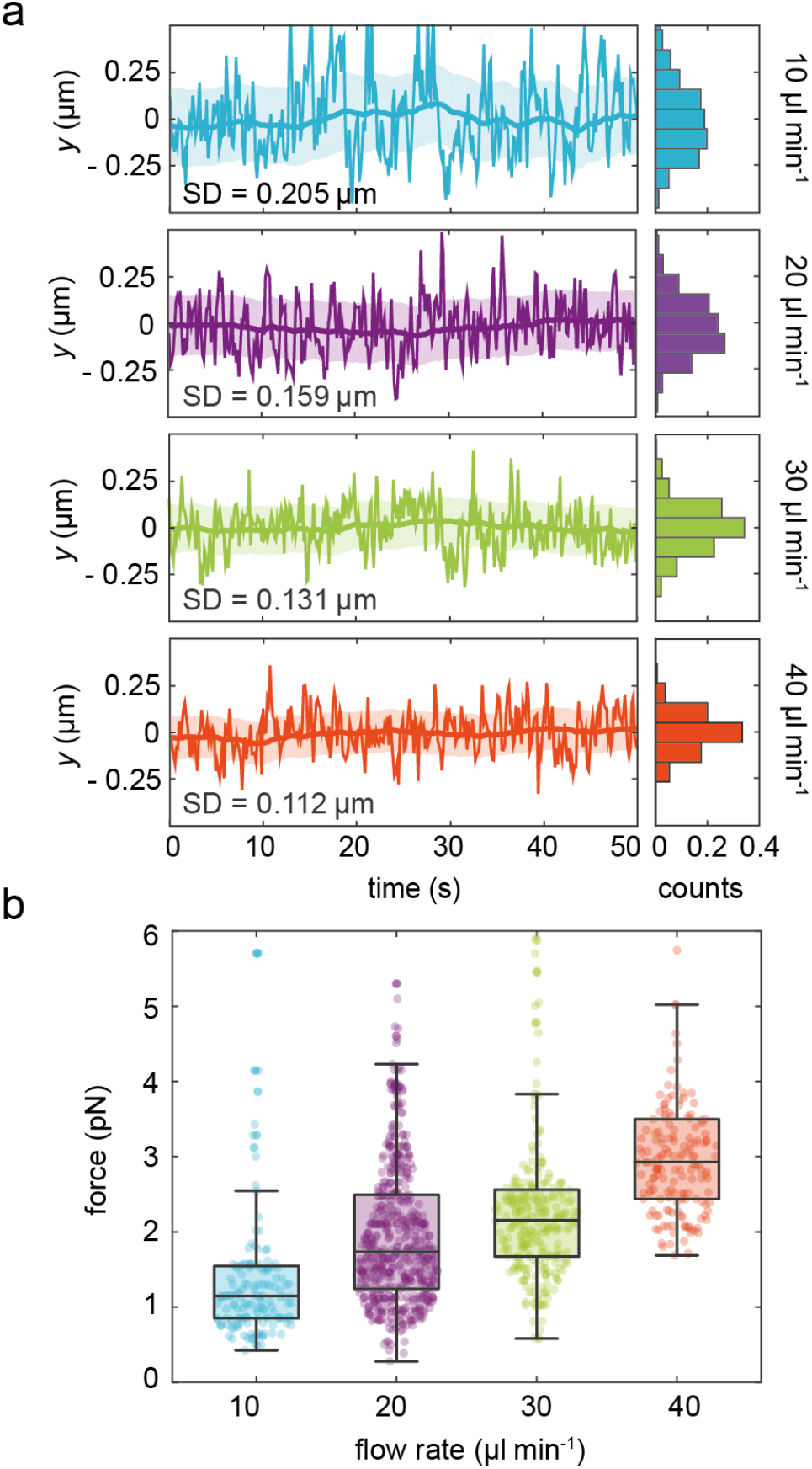
Fluctuation-based in situ force calibration. a) Bead fluctuations in the direction *y* transverse to the flow over time for flow rates of 10, 20, 30 and 40 μl min^−1^. Histograms on the right-hand side present relative occurrences of *y* positions. b) Distributions of estimated forces for the different flow rates from a single experiment performed at a given flow rate. The forces were estimated based on the bead fluctuations displayed in a. Boxes extend from 25th to 75th percentiles, with a line at the median. Whiskers span 1.5 × interquartile range. Colored dots represent individual beads (*n* = 162, 553, 285, 169).

### 2.4. Motility of individual kinesin-1 motors under resisting and assisting loads

After AMPPNP had been washed out by ATP-containing buffer the motors started to translocate (**Figure S1**), predominantly along the flow axis as the majority of the microtubules were aligned by the flow (**Figure S3**). We discriminated between different stepping directions by looking at the bead displacement in the *x*-*y* plane. Exemplary trajectories of kinesin-1 motors moving against the flow (i.e. experiencing a resisting load) and with the flow (i.e. experiencing an assisting load) are presented in **Figure 3a** and **3b**, respectively.

**Figure 3.**
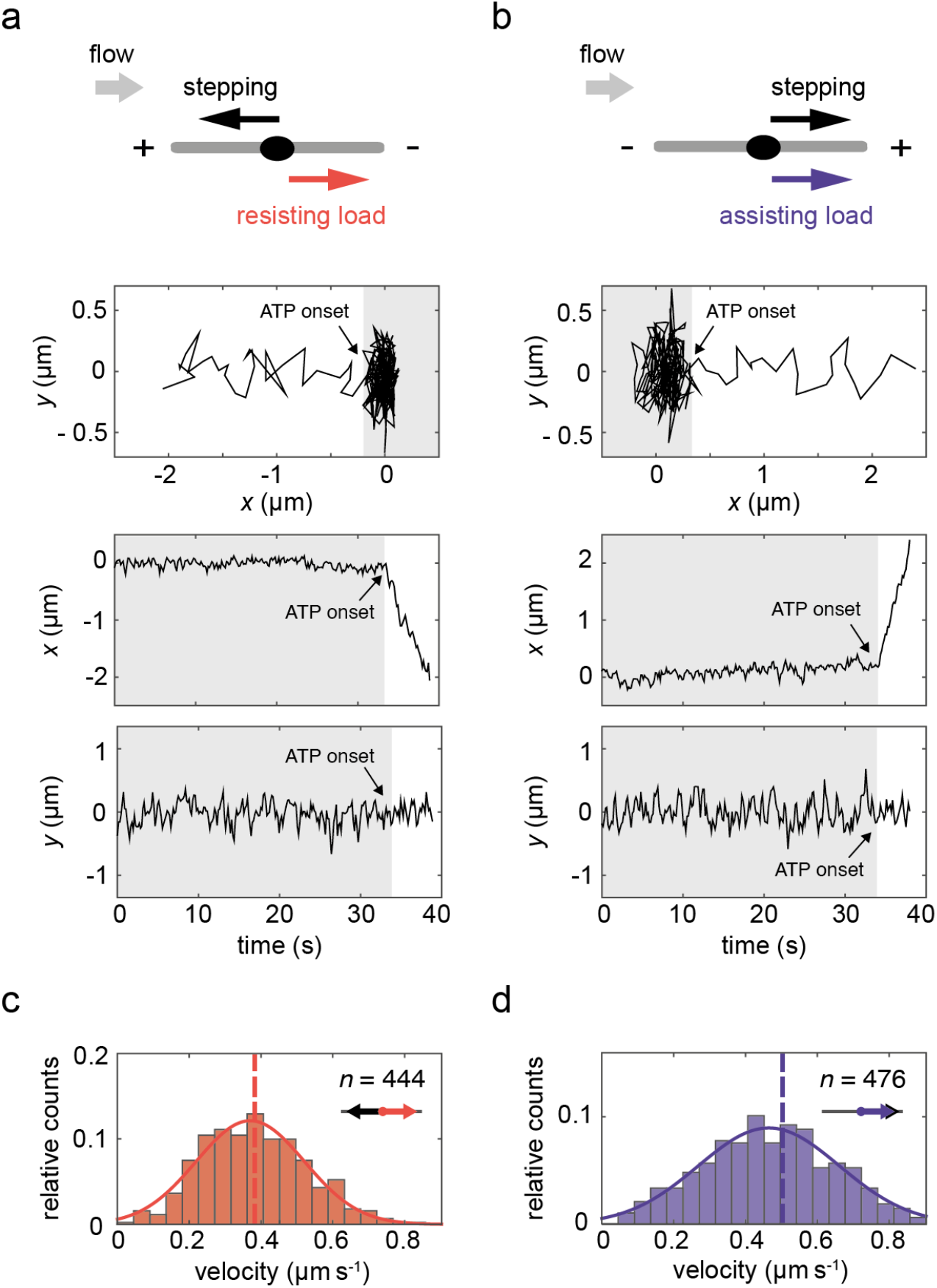
Single-molecule motility events under resisting and assisting load. a) Setup and exemplary trajectory of a motor stepping against the flow (towards decreasing *x*-position, corresponding to a resisting load). The plots from top to bottom present: the position of the bead in a 2D plane over 40 seconds before detachment, the *x*-position over time and *y*-position over time. The gray-shaded areas correspond to the period before ATP onset, i.e. the time when the motors were still arrested in the presence of AMPPNP. b) Analogous setup and exemplary trajectory of a motor stepping with the flow (towards increasing *x* position, corresponding to an assisting load). c) Velocity histogram of 444 stepping events under resisting load. d) Velocity histogram of 476 stepping events under assisting load. In c and d, overlays of Gaussian fits are presented and vertical dashed lines represent mean velocity values, the histograms represent events recorded in one experiment performed under 20 μl min^−1^ flow rate.

Although the microtubule axes were mostly aligned with the flow direction, their polarities (i.e. the positions of their plus and minus ends) were arbitrary (**Figure S3**). Therefore, we were able to investigate the motility of individual plus-end directed kinesin-1 motors under resisting and assisting loads of the same magnitude simultaneously. Velocity histograms from a single experiment with 444 motility events against the flow and 476 motility events with the flow are presented in **Figure 3c** and **3d**. Under a median load of 1.7 pN in the presented experiment, the kinesin-1 motors stepped with mean velocities of 0.424 ± 0.008 μm s^−1^ against the flow and 0.557 ± 0.014 μm s^−1^ (mean ± SEM) with the flow.

### 2.5. Kinesin-1 force-velocity curve

By varying the flow rates, we applied loads of different magnitudes and constructed a force-velocity curve for kinesin-1 (**Figure 4a**). Since the distribution of forces acting on the molecules under a given flow rate is considerably broad (**Figure 2b**), we assigned force loads to each stepping event individually and compared velocities for the data grouped according to the estimated values into 0.3-pN wide bins. We observe that with increasing resisting load the stepping velocity of kinesin-1 progressively decreased. It reached a mean value of 0.510 ± 0.031 μm s^−1^ (mean ± SEM) for the lowest load bin (0.6 ± 0.15 pN) and 0.250 ± 0.016 μm s^−1^ for the highest load bin (3.3 ± 0.15 pN). Under conditions of assisting load, we found the highest velocity of 0.703 ± 0.050 μm s^−1^ for the lowest load bin. With increasing assisting load, the kinesin-1 stepping velocity slightly decreased and reached a mean value of 0.431 ± 0.034 μm s^−1^ for the highest load bin.

**Figure 4.**
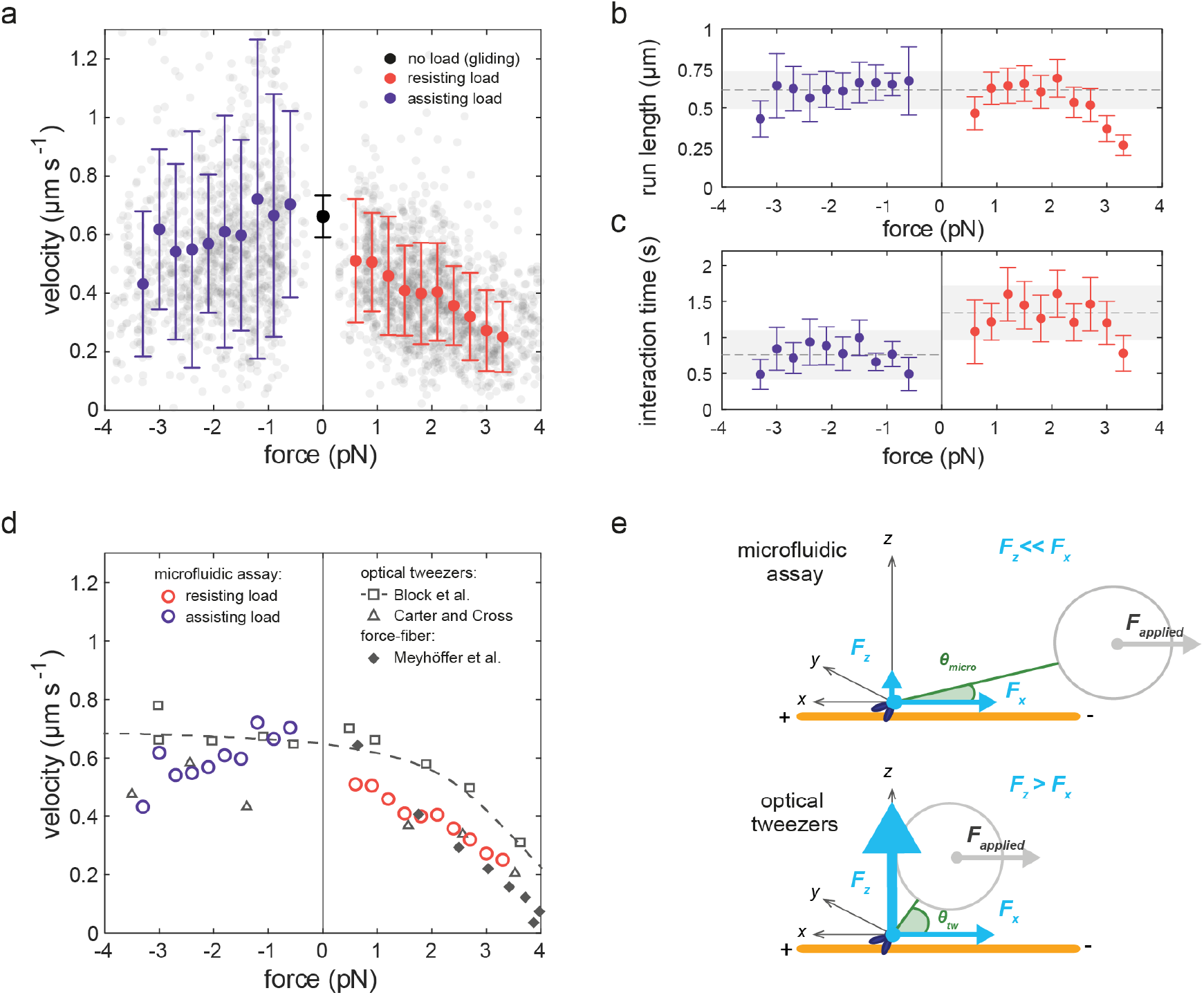
Force-dependence of motility parameters for kinesin-1 as probed by the microfluidic assay. a) Stepping velocity of kinesin-1 observed under assisting (violet dots) and resisting (red dots) loads of different magnitudes. Plotted values represent means ± SD of velocity data contained within 0.3-pN force bin (*n* = 52–213 per bin). Gray scatter represents individual events used for binning (*n* = 2499 events, out of which 2170 are included in the force bins, data pooled from eight independent experiments performed at 10–50 μl min^−1^). The black dot represents the velocity of the kinesin-1 motor used in this study under no load condition as evaluated by gliding assays (mean ± SD, see **Figure S4**). b,c) Force dependence of run lengths and interaction times for events presented in (a), plotted values represent means ± 2 × SD obtained from bootstrapping. As guides for the eye, means of bins −3 to 2.7 pN for run length, and mean of all assisting load bins, as well as all resisting load bins up to 3 pN for interaction time, are indicated with the dashed gray line. Shaded areas represent 2 × SD. d) Overlay of force-velocity data from our study (open circles) with data obtained in optical tweezers’ studies by Block et al.^[37]^ (open squares and dashed curve showing the fit of a five-step model) and by Carter and Cross^[38]^ (open triangles), and in a force-fiber assay by Meyhöfer et al.^[22]^ (filled diamonds). e) Comparison of experimental geometries in the microfluidic assay presented in this study (upper scheme) and in a conventional optical tweezers experiment (lower scheme). *θ* - inclination angle between the microtubule and the tether in the case of the microfluidic setup (*θ*_*micro*_) and a conventional optical tweezer configuration (*θ*_*tw*_); *F*_*applied*_ – force applied on the bead; *F*_*z*_ – vectorial component of the force pointing in the *z* direction (vertical force); *F*_*x*_ – vectorial component of the force pointing in the *x* direction (horizontal force).

The force-velocity curve obtained for kinesin-1 in our study follows qualitatively^[37]^ and even quantitatively^[38]^ earlier data from optical tweezers, as well as from force-fiber^[22]^ measurements for resisting loads (**Figure 4d**). We note, that contrary to the data reported in by Block et al.^[37]^ but consistent with the trend reported by Carter and Cross^[38]^ the velocity observed in our experiments showed a marked decrease under increasing assisting loads. The decreased velocity of kinesin-1 under assisting loads could potentially be explained by sterical hinderence caused by the motor stalk being drawn towards the microtubule surface in front of the motor head.

### 2.6. Dependence of further motility parameters on applied force

Apart from velocity, we were also able to readily evaluate the run lengths and interaction times of individual kinesin-1 motors under the applied loads (**Figure 4b–c**). To account for under-representation of very short stepping events, we estimated these two parameters using least-squares fitting of the cumulative distribution function with free cut-off parameters.^[44]^ The measured run lengths appeared fairly constant for loads between −3 pN and 2.7 pN, with a mean at 0.63 μm (**Figure 4b**). For assisting loads larger than 3 pN and resisting loads larger than

2.7 pN the run lengths decreased. This is in contrast to observations made in conventional optical tweezers experiments, where the measured run lengths decreased already drastically for moderate loads, e.g. showing a three-fold decrease at 2 pN resisting load.^[45]^ This discrepancy can be attributed to the high vertical forces in conventional optical tweezers experiments that cause pre-mature detachment of motors compared to forces applied more horizontally.^[46–48]^ The mean run-length value of 0.63 μm observed at low loads, corresponds well to the previously observed value of 0.67 μm for unloaded motors.^[44]^

The measured interaction times appeared fairly constant for all applied assisting loads, with a mean at 0.81 s (**Figure 4c**). For resisting loads the interaction times appeared constant up to 3 pN, with a mean at 1.43 s, and decreased above that load. The overall higher interaction times observed under resisting loads, as compared to assisting loads, suggest that kinesin-1 exhibits a higher detachment rate under assisting loads. This is in agreement with the higher unbinding force observed for kinesin-1 under resisting load as compared to assisting load^[49]^ and with the theoretical prediction that horizontal forces alone, as predominantly present in our setup, decelerate motor detachment.^[46,47]^ The mean interaction times obtained for both assisting and resisting loads, correspond well to the interaction time of 0.95 s under unloaded conditions reported previously for the kinesin-1 at room temperature.^[44]^ The good agreements of both run lengths and interaction times with previously reported and predicted values for single kinesin-1 motors ascertains that we evaluated the stepping of single molecules.

### 2.7. Reducing the vertical force component

Our hydrodynamic force assay not only enables parallelization of the measurements on cytoskeletal motors, but also provides an alternative geometry of force application compared to existing methods. While optical traps—the method of choice for characterizing cytoskeletal motors—have been exploited to study the application of forward, backward,^[38,45,50]^ and sideward loads^[37,51]^ on stepping kinesins using a variety of geometries,^[52,53]^ they generally suffer from a poor control over vertically applied forces, which may bias the measurements performed.^[23,25,26,54]^ Although the force is applied horizontally onto the bead in a conventional optical trap experiment, the molecule under investigation is experiencing a vertical load which, in fact, can surpass the applied horizontal force in magnitude.^[25]^ Such substantial vertical load is pulling the motor away from the filament and can influence its detachment rate.^[23,25,46–48]^ In our approach, the introduction of a spacer between the motor and the bead reduces the vertical force component to less than 15% of the applied force (see **Figure 4e** and **Table S1**), thus applying forces more stringently in the direction of motor movement. Such a prolonged spacer could, in principle, be implemented in conventional optical tweezer experiments. A reduction of vertical force component in optical tweezers can also be achieved in a three-bead assay with suspended microtubules, as recently demonstrated.^[48]^ To explore the influence of loading geometry on the measured motility parameters in our assay, experiments with varying linker length or varying vertical magnetic force could be performed. The different loading scenarios may reflect the physiological transport of cargos differing in size and shape inside the cell, or differently positioned motors in multi-motor assemblies present in vivo.

## 3. Discussion

The high-throughput microfluidic assay devised in this work presents a platform for multiplexed studies of cytoskeletal motor protein behavior under external loads. Leveraging the large spatially homogenous force field generated by hydrodynamic flow and the use of a specialized telecentric lens capturing a field of view of several millimeters in size, we were able to optically track hundreds to thousands of individual molecules in a single experiment. To demonstrate the power of the approach, we studied the dependence of the motility parameters on the applied load for the microtubule-based motor protein kinesin-1, the major workhorse of cargo transport in living cells. We applied a series of in situ calibrated loads and observed that the velocity of kinesin-1 decreased under resisting and assisting loads by up to 62% and 35%, respectively, at the maximum applied load of 3.3 pN. The number of evaluated molecules amounted to roughly 2500 in eight experiments, enabling hydrodynamic force–spectroscopy measurements on cytoskeletal motor proteins at unprecedented throughputs.

### 3.1. Overcoming the multiplexing challenge for the case of cytoskeletal motors

Parallelization of the force application for single-molecule studies is a challenging endeavor and is especially hard to accomplish for the case of cytoskeletal motors. Multiplexing with optical tweezers is generally possible, but the number of beads that can be held by constant and appreciable forces is rather low due to the splitting of the finite total optical force.^[6–8]^ Magnetic tweezers, on the other hand, are capable of applying forces to multiple molecules at a time,^[55,56]^ however, potential variations of the applied force field with the vertical and horizontal position must be taken into consideration.^[57]^ To overcome these limitations, we implemented hydrodynamic flow to create a large-area, homogenous force field and a microscopy setup optimized for large fields of views, as *x-y* position tracking for beads with large scattering cross-section can provide satisfactory accuracy at low-magnifications. The use of the recently developed multiplexed single-molecule methods such as acoustic force spectroscopy^[16]^ and centrifuge force miscrosopy^[15]^ have been demonstrated on several systems. However, their application to cytoskeletal motors is not straightforward, due to the vertical character of the applied force.

### 3.2. Method performance and additional assets

Optimal performance of our method is achieved at intermediate forces. At very low forces (< 0.5 pN) the bead fluctuations limit the accuracy of the velocity measurements. At high forces, in turn, the observation time is limited due to the decreased processivity of the motor. However, the latter is specific to single kinesin-1 motors, exhibiting a force-dependent run length,^[45]^ and will likely not be an issue for other mechanosystems, such as dynein^[58,59]^ or multi-motor transport systems.^[60]^

An additional asset of our method is the possibility to study the motor velocity in an angle-resolved manner (see **Figure S5**). Depending on the alignment of the microtubules with respect to the flow direction, some motors will step not directly against or with the direction of applied force. In this study, the microtubules were aligned with the long flow cell axis to maximize the number of events stepping parallel to the force direction (**Figure S3**), however, the orientation of the microtubules can be randomized by applying a perpendicular or turbulent flow while introducing microtubules to the flow cell.

The straightforward in situ force calibration and the flexibility of the assay geometry further enhance the appeal of the presented method. In terms of geometry, the angle at which the force is applied to the motors can be easily adapted by changing the tether length or by adjusting the magnitude of the magnetic force that lifts the beads. Additionally, similarly to previous studies using DNA origami^[61]^ or flexible DNA scaffolds^[62]^, the DNA tether could be engineered to accommodate multiple attachment sites for motors. This would offer a possibility to study processivity and speed of multi-motor assemblies, that are relevant to physiological scenarios, under force. Finally, we note that our approach can be implemented using any standard wide-field microscope at low cost.

## 4. Conclusion

Single-molecule manipulation techniques have shed light on the functioning principles of many molecular machines in the cell. Yet, their widespread applicability and utilization for single-molecule screening purposes is limited by the lack of robust high-throughput technologies. Our versatile, massively multiplexed microfluidic assay for the application of forces to molecular mechanosystems, such as stepping cytoskeletal motors and motor complexes, presents a leap towards wider usage of single-molecule approaches. We envision a broad implementation of the assay in fundamental research of biological systems as well as in medical diagnostics applications, where rapid acquisition of population-wide data is of key importance.

## 5. Methods

### Protein production and purification

We used a truncated kinesin-1 heavy chain from *Rattus norvegicus* (1-430 aa) fused to a SNAP-tag and 8xHis-tag (rKin430-SNAP-8xHis in a pET17b, see **Figure S6** for full protein sequence). The SNAP-tag allows for covalent binding of O^6^-benzylguanine (BG) to the protein.^[63]^ Protein expression was performed in *Escherichia coli* BL21(DE3) pRARE (Invitrogen) under T7 promoter with 1 mM isopropyl-β-D-thiogalactopyranoside (IPTG) induction at OD600 = 0.6 for 14 h at 18 °C. After protein expression, bacterial cells were disrupted in a high-pressure homogenizer (Emulsiflex-C3, Avestin Inc) in the presence of protease inhibitors (Protease Inhibitor Cocktail Tablets, Roche Diagnostics GmbH). Crude lysate was centrifuged at 20,000g at 4°C. The supernatant was loaded on a HisTrapTM metal affinity column (GE Healthcare Life Sciences). All protein purification steps were performed in buffers based on 2 x phosphate-buffered saline (PBS; 274 mM NaCl, 5.4 mM KCl, 16.2 mM Na_2_HPO_4_ 2H_2_O, 3.52 mM KH_2_PO_4_, 2 mM MgCl_2_, pH 7.4) containing 1 mM ATP and 1 mM dithiothreitol (DTT). Column washing was performed with 10 times the column volume of: 5 mM ATP in 2 x PBS, 6 x PBS, 15 mM and 30 mM imidazole in 2 x PBS; followed by elution with 500 mM imidazole in 2 x PBS. Size and purity of the protein after elution was checked by SDS–PAGE (**Figure S7**). The peak fractions were then desalted on size-exclusion Sephadex columns (NAP25 gravity column, GE Healthcare). Collected protein was snap-frozen in liquid nitrogen and stored at −80°C. To check the activity of the obtained protein, gliding motility assays were performed with specific immobilization of motors on the surface via penta-His antibodies (34660, Qiagen) as previously described^[64]^ (**Figure S4**). Microtubule movement was tracked using FIESTA software.^[65]^ On basis of Gaussian fitting the mean gliding velocity for KinSNAP was estimated to be 662 ± 72 nm s^−1^ (mean ± SD; *n* = 243). This velocity corresponds well to literature data as the microtubule gliding velocity at saturating ATP concentrations and pH 6.9 is reported to fall between 500 and 750 nm s^−1^.^[66]^

### Microtubule polymerization

Microtubules stabilized with GMPCPP (Jena Bioscience, Germany) were prepared by polymerization of in-house prepared porcine tubulin labelled partially with rhodamine (1:3 ratio of labelled to unlabeled tubulin). Polymerization was carried out using 0.25 mg ml^-1^ tubulin in BRB80 buffer (80 mM piperazine-N,N’-bis(2-ethanesulfonic acid) (PIPES), 1 mM MgCl_2_, 1 mM ethylene glycol tetraacetic acid (EGTA), pH 6.9 adjusted with KOH) supplemented with additional MgCl_2_ to final concentration of 4 mM and 1 mM GMPCPP. The polymerization reaction was pre-incubated for 10 min on ice and continued for 2 h at 37°C. Afterwards, to remove unpolymerized tubulin dimers from the solution, the microtubules were spun down in a tabletop microcentrifuge (Heraeus Fresco 17, Thermo Scientific Inc.) at 17000 g for 15 minutes and resuspended in 250 μl of BRB80 buffer.

### Preparation of doubly-functionalized DNA linkers

λ-DNA (N3011, NEB) was functionalized on one end with *O*^*6*^-benzylguanine (BG) and on the other with digoxigenin (Dig) by two-step ligation of oligonucleotides with functional groups to the λ-DNA overhangs. The Dig-oligonucleotide was purchased in functionalized form (AGGTCGCCGCCCA_12_-Dig, Eurofins MWG Operon). The BG-oligonucleotide was prepared by coupling of 10 mM BG–GLA–NHS (S9151, NEB) to 0.33 mM amine-functionalized oligonucleotide (GGGCGGCGACCT-NH_2_, Eurofins MWG Operon). The coupling reaction was conducted in 67 mM HEPES (pH 8.5) and 50% DMSO for 30 min at room temperature. The uncoupled BG-GLA-NHS was removed by filtration on Micro Bio-Spin™ P-6 Gel Columns (Bio-Rad).

The BG-oligonucleotide (210 nM) was ligated to the 3’ end of the λ-DNA (3 nM) in 500 μl T4 ligase buffer (50 mM Tris-HCl, 10 mM MgCl_2_, 1 mM ATP and 10 mM DTT, pH 7.5, NEB). Before adding the ligase, the solution was incubated for 5 min at 65°C and cooled down slowly to allow for annealing. For the ligation 800 units of T4 DNA ligase (NEB) were added and the reaction was held overnight at room temperature. Next, Dig-oligonucleotide (460 nM, a 3-fold excess with respect to BG-oligonucleotide) was annealed to the 5’ end of the λ-DNA by incubating at 45°C for 30 min. After cooling the solution down to room temperature, additional 800 units of T4 DNA ligase were supplemented and the ligation reaction was held for 2 h at room temperature. To get rid of the remaining oligonucleotides and oligonucleotide duplexes, the ligation product was dialyzed at 4°C against 0.5 l TE buffer (50 mM Tris-HCl, 1 mM EDTA) using 1 ml Float-A-Lyzer G2 Dialysis Device with molecular cutoff of 1000 kDa (G235037, Spectrum Labs). The dialysis buffer was exchanged 5 times with 4–16 h intervals.

### Bead functionalization

Superparamagnetic polystyrene beads with a diameter of 1.08 μm (coefficient of variation < 5%, Dynabeads MyOne™ Tosylactivated, 65501, Invitrogen) were functionalized via an amine coupling reaction with 20 μg of anti-digoxigenin Fab fragments (Invitrogen) per mg of beads according to a protocol described elsewhere.^[67]^ Of note, the size of the beads chosen for the assay will determine the force applied to the motors at a given flow rate and influence some practical aspects of the experiment such as the efficiency of forming functional tethers or the maximal motor density in case of a dense surface coverage. We experienced that 1-μm beads are more efficient in forming tethers than 2.8-μm beads from the same provider. 1-μm beads also offer a lower sedimentation rate what reduces the number of beads stuck to the surface during assay preparation that cannot be easily lifted later on.

### Coverslip functionalization

For making their surface hydrophobic, the coverslips used for all experiments were coated with DDS (dichlorodimethylsilane) prior to use. DDS-functionalization involves several cleaning steps, including 60 min incubation in Piranha solution (30% H_2_O_2_ and 70% H_2_SO_4_) and a silanization step using DDS diluted in TCE (trichloroethylene). Details can be found elsewhere.^[68]^

### Flow cell assembly

For performing the microfluidic assays, a PDMS slab with 3 mm wide and 100 μm high channel was placed on the 24 × 60 mm DDS-functionalized coverslip and pressed with a custom-made metal frame to avoid leakage. Flow channels terminate with Y junctions at each end providing for two inlets and two outlets (**Figure S8**). The solutions were introduced to the flow cell by a pump (AL-200, WPI, Inc) operated in withdrawal mode through polyethylene tubing (PE-60, 0.76 mm inner and 1.22 mm outer diameter) inserted into 1.2 mm holes punched into the PDMS. Volumetric flow rates of 10, 20, 30, 40 or 50 μl min^−1^ (corresponding to calculated average flow velocities of 0.6, 1.1, 1.7, 2.2 or 2.8 mm s^−1^) were kept constant during the measurements. For the flow cell dimensions and flow velocities used the flow is laminar (*Re* < 1, see **Table S2**).

### On-surface assay assembly

Anti-β-tubulin antibodies (75 μg ml^−1^, SAP.4G5, Sigma-Aldrich) in BRB80 was flushed into the flow cell and incubated for 5 min to allow for hydrophobic interaction-based absorption to the surface. Next, the channel surface was passivated with 1% Pluronic F127 (P2443, Sigma-Aldrich) in BRB80 for 30-60 min. The flow cell was then mounted on the imaging setup and connected to the syringe pump operated at 20 μl min^−1^ throughout the following assembly steps. First, the flow cell was washed with 200 μl of BRB80 solution. Next, 250 μl of GMPCPP-stabilized microtubule solution in BRB80 was flowed through the channel. SNAP-tagged kinesin-1 (11 nM) was pre-coupled to the functionalized DNA linker (0.4 nM) by incubation for 2 to 3 h at room temperature on a rotary mixer and diluted 1:10 in imaging solution (0.2 mg ml^−1^ casein, 10 mM DDT, 0.1% Tween 20 in BRB80) containing 100 μM AMPPNP. 200 μl of so-prepared kinesin-DNA complexes was introduced to the channel. After a subsequent wash with 200 μl of imaging solution, anti-digoxigenin beads were flowed through the flow cell and attached on the fly to the kinesin-bound DNA linkers. Subsequently, another washing step was performed to remove the unbound beads from the flow cell. During this step the flow rate was adjusted to the desired value (between 10 and 50 μl min^−1^) and a magnet was placed at a defined height above the flow cell to minimize interaction of the tethered beads with the surface. The magnet position was controlled by a metal arm with a built-in ruler. The applied magnetic force was estimated to be 0.1 pN using the bead fluctuations at no flow condition, as established for magnetic tweezers^[41]^. Finally, to initiate kinesin stepping, an imaging solution containing 10 mM ATP was flowed into the flow cell at the respective flow rate.

### Imaging setup and data acquisition

A fiber illuminator (Thorlabs) was used to illuminate the flow cell from the side. The light scattered by the beads was collected through a telocentric lens (TL12K-70-15, Lensation) with 7 × magnification mounted directly on top of a 29 Megapixel CCD camera (Prosilica GX6600, Allied Vision Technologies, 5.5 μm pixel size). Images were collected at 150 ms per frame continuously in streaming mode using StreamPix imaging software (NorPix). All the experiments were performed at room temperature (∼23°C).

### Data analysis

The centroid of the beads was tracked using a custom ImageJ plugin programmed in house, the core of the tracking algorithm corresponds to the *Peak Tracker* (https://duderstadt-lab.github.io/mars-docs/docs/image/PeakTracker/) implemented in the open-source **M**olecular **A**rchive **S**uite, *mars*, software (https://github.com/duderstadt-lab/mars-core). Bead trajectories were corrected for drift by subtracting an averaged trace of several immobile particles. For the velocity calculations, the distance along the line fitted to the *x-y* displacement of the bead and time were used. The start and end times of the stepping events were marked by hand.

### Localization precision

To determine the localization precision for bead tracking, we estimated the standard deviation of *x* and *y* positions for 12 beads stuck to the surface over the time of 5 min. The used traces were corrected for drift by subtracting the average of all stuck bead traces. The obtained standard deviations averaged at 32 nm for both *x* and *y*. This value corresponds to the experimental value of localization precision, the theoretical limit of the tracking precision in the used configuration was previously calculated to be 6 nm.^[36]^

### Estimation of the acting force

In our system a bead is tethered with a dsDNA tether with extension length *l* to the kinesin-1 motor attached to the microtubules on a glass slide (**Figure S9a**). The fluctuations of the bead in the direction perpendicular to the flow, *y*, can be approximated by the movement of a pendulum with a length corresponding to the extension of the tether *l* (**Figure S9b**,**c**). During the stochastic movement of the bead, the force *F*_*spring*_ is pushing it back to the equilibrium state and is defined as:

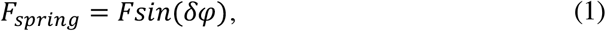

where *F* is the applied drag force and *δφ* is the angle between *F* and the tether (**Figure S9c**).

For small angles *sin*(*δφ*), can be approximated as 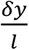 and the above equation takes form:

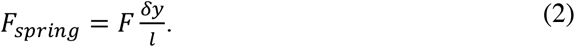

Therefore, in our Hookean system the spring constant defining the stiffness is described as:

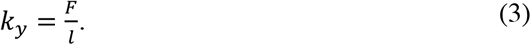

The mean energy of a Hookean system is expressed as follows:

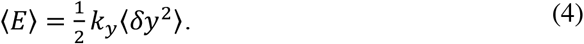

Relating this equation to the equipartition theorem yields:

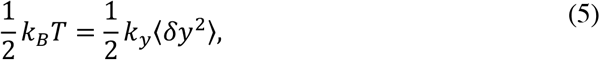

where *k*_*B*_ is the Boltzmann constant and *T* is the temperature. We can therefore relate the force *F* to the root-mean-square deviation of position in *y* in the following way:

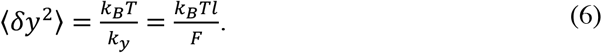

Similar approaches were used for force estimation in previous studies using magnetic tweezers,^[41]^ as well as microfluidic DNA stretching.^[31]^

For estimating the bead fluctuations, only trajectories with subsequent stepping events were considered. For each trajectory a fragment of a minimum of 100 data points of the position in the y direction was chosen. These fragments were visually inspected to discard trajectory regions where abrupt position changes or loss of fluctuations due to sticking of the bead to the surface were present. The measured mean square displacement of the bead position in y, ⟨*δy*^2^⟩_*m*_, was estimated according to the formula:

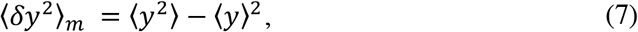

and corrected for motion blurring caused by the finite camera integration time *W*, in our case equal to 150 ms, using a correction function *S*(*α*)^[69]^ as follows:

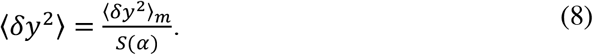

The correction function *S*(*α*) is given by^[69]^:

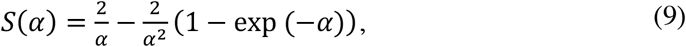

where *α* is the ratio of the camera integration time *W* to the trap relaxation time *τ*

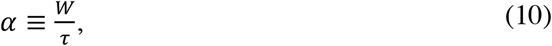

with 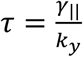, where *k*_*y*_ is the spring constant of the trap introduced in Equation (3), and the Stoke’s friction coefficient corrected for the surface proximity using a correction factor dependent on the *z* position for bead movement parallel to the surface λ _| |_ (*z*) derived by Faxen^[70]^ (see **Supplementary Note 1** for more details), γ _| |_ = λ_| |_ (*z*) *γ* = λ_| |_ (*z*) 6 π η*R*, where η represents dynamic viscosity of the medium and the radius of the bead. Hence, *α* can be expressed as:

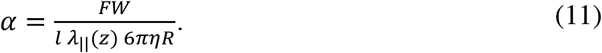

Since *W*, η, and *R* are constant and dictated by the experimental conditions, *α* becomes a function of the applied drag force *F*, the extension of the tether *l*, and the correction factor λ _| |_ (z).

Taking into account the motion blurring correction, **Equation 6** can be rewritten as:

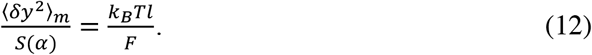

Since the tether extension *l* is not verified experimentally in our assay, we cannot estimate force using **Equation 12** alone. We therefore took advantage of the force-extension relation that describes the behavior of a DNA strand when pulled by a force^[42]^:

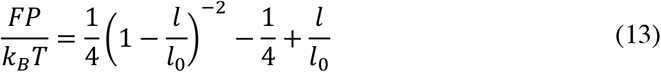

with *l*_0_ corresponding to the contour length of the λ-DNA (16.2 μm) and to the persistence length of double stranded DNA (50 nm)^[42]^, to create a set of equations (**Equation 12** and **13**). The obtained set of two polynomial equations was solved numerically (using *vpasolve* function in MATLAB R2020a, The MathWorks, Inc., Natick, MA, USA) to obtain the values of *F* and *l* for each measured molecule.

### Estimation of interaction times and run lengths

Run lengths and interaction times were estimated using least-squares fitting of the following cumulative distribution function:

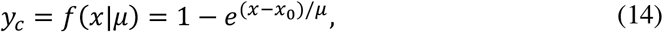

with *x* epresenting the measured parameter (run length or interaction time), μ the population mean, and *x*_0_ a free-fitted cut-off parameter introduced to account for underrepresentation of very short events^[44]^. For estimating the mean and standard deviation values of the measured parameters the bootstrapping approach was used, in that the distributions were resampled with replacement 100 times and fitted each time. The means and standard deviations of the μ values fitted in all resampling rounds are reported.

## Data Availability Statement

Source data for the figures are provided as a supplementary file alongside the manuscript. The datasets containing tracking results supporting findings of this study, together with analysis scripts, are available on figshare (http://www.doi.org/10.6084/m9.figshare.14071382).^[71]^ Post-processed part of the data accompanied by the analysis scripts is also available on GitHub (https://github.com/MartaUrb/microKBA).

## Author Contributions

S.D., A.M.O. and W.J.W. conceived the project. M.U., with support of A.L., W.J.W. and K.E.D. performed the experiments. W.J.W. cloned and expressed the SNAP-tagged kinesin-1 protein. M.U. and K.E.D. wrote the analysis software. M.U. analyzed the data, prepared figures and wrote the original version of the manuscript. All authors revised and edited the manuscript. S.D., K.E.D., A.M.O. and W.J.W. supervised the project. S.D. and K.E.D. acquired funding.

## Acknowledgement

We thank Corina Bräuer and Cornelia Thodte for technical assistance, Felix Ruhnow, Erik Schäffer and Joe Howard for fruitful discussions, Lucas Wittwer and Sebastian Aland for help with theoretical considerations regarding bead position, and Georg Krainer for helpful suggestions on the manuscript. We acknowledge financial support from the Dresden International Graduate School for Biomedicine and Bioengineering (DIGS-BB, stipend to A.L.), the German Excellence Initiative (TU Dresden Support-the-Best grant to S.D.), the Human Frontier Science Program (Long-term fellowship LT000180/2012-L to K.E.D.), and support from the Max Planck Society to K.E.D.

