## Supplementary Information for "Highly-parallel microfluidics-based force spectroscopy on single cytoskeletal motors"

bioRχiv, https://doi.org/10.1101/2020.08.11.245910

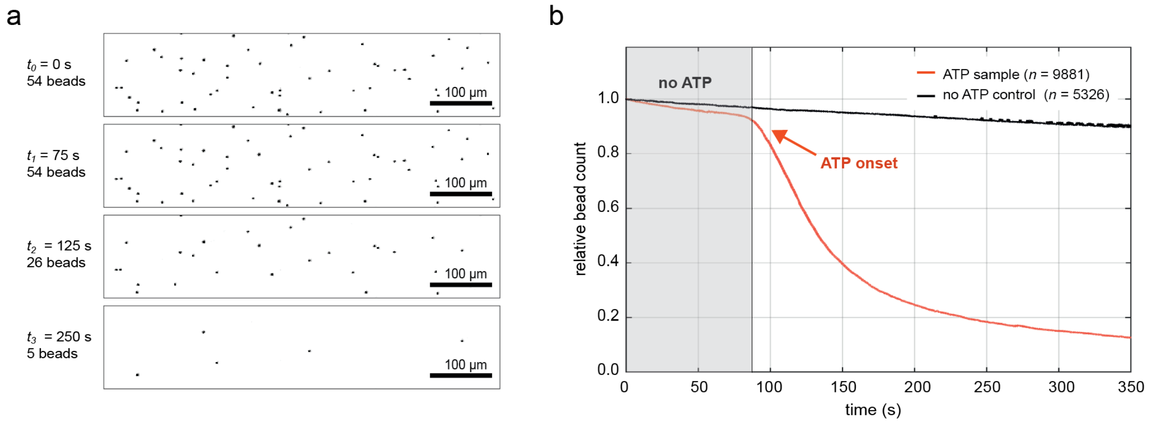


**Figure S1**. Onset of kinesin-1 motility upon ATP delivery. Detachment of the kineisn-bead constructs (due to the limited run length of kinesin-1) in presence of ATP constitutes a global footprint of the kinesin-1 enzymatic activity. The molecules are first immobilized on microtubules with 100 µM AMPPNP ($t_0$) and gradually induced to step and detach after addition of 10 mM ATP to the flow cell at around $t$ = 90 s. a) Images of 0.5 % of the total field of view visualizing a decreasing number of beads over time. b) Relative bead count over time in an exemplary experiment when ATP was introduced (red curve) as compared to a control experiment where no ATP was added (black curve). Total number of beads detected at $t_0$ equals 9881 and 5326 for ATP and no ATP samples, respectively.

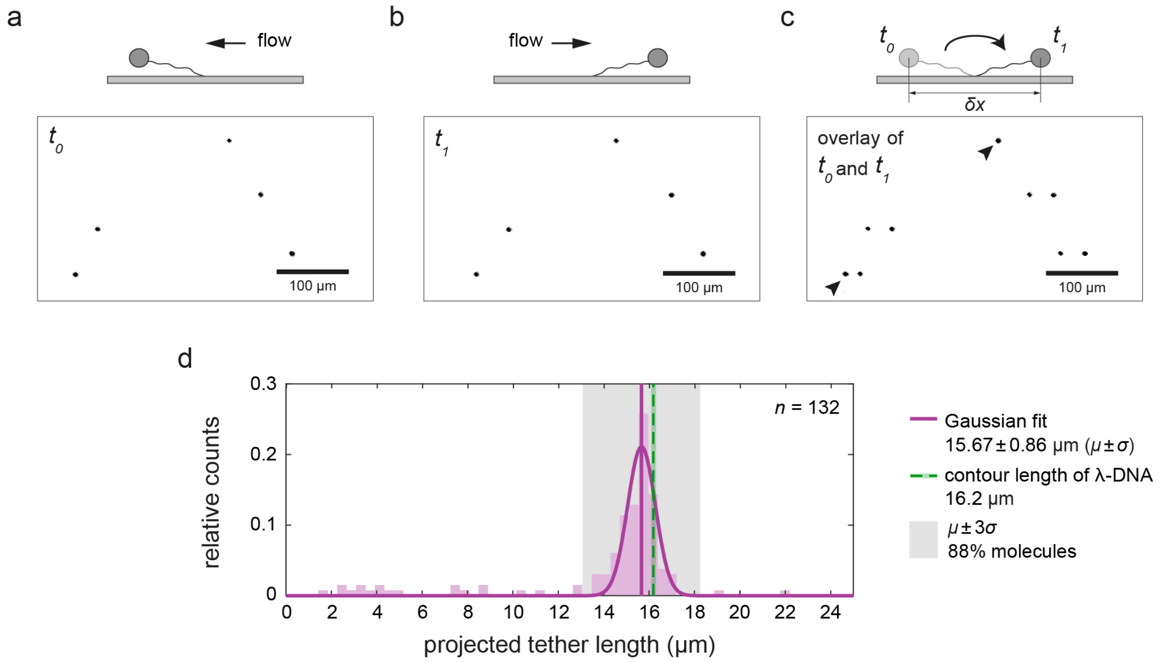


**Figure S2.** Majority of tethered beads show close to full-length tether extension during flow reversal. a) Fraction of the field of view showing bead positions at time $t_0$, when flow was applied in the direction from right to left. b) Same fraction of the field of view time $t_1$, when flow direction was changed to come from left to right. c) Overlay of a and b. The three beads in the middle correspond to full length tethers. The top bead (indicated by an arrowhead) did not change its position over time and corresponds to a bead stuck to the surface. The bead in the lower left corner (indicated by another arrowhead) shows a position difference smaller than full extension of the tether and may correspond to a bead with entangled tether. d) Histogram of projected tether lengths for beads that changed the position during flow reversal ($n$ = 132). 88% of measured tethers reached values within 3 standard deviations from the Gaussian peak (shaded in gray), corresponding to full-length, unentangled tethers. Projected tether lengths where calculated as follows $\frac{\delta x - 2R_{bead}}{2}$, where $R_{bead}$ corresponds to the bead radius, and $\delta x$ to the bead-to-bead distance measured in *Fiji* by drawing a line between the two extreme bead positions on the overlay image. The contour length of the $\lambda$-DNA, corresponding to maximal possible extension of the tether, is plotted as a green dashed vertical line for reference.

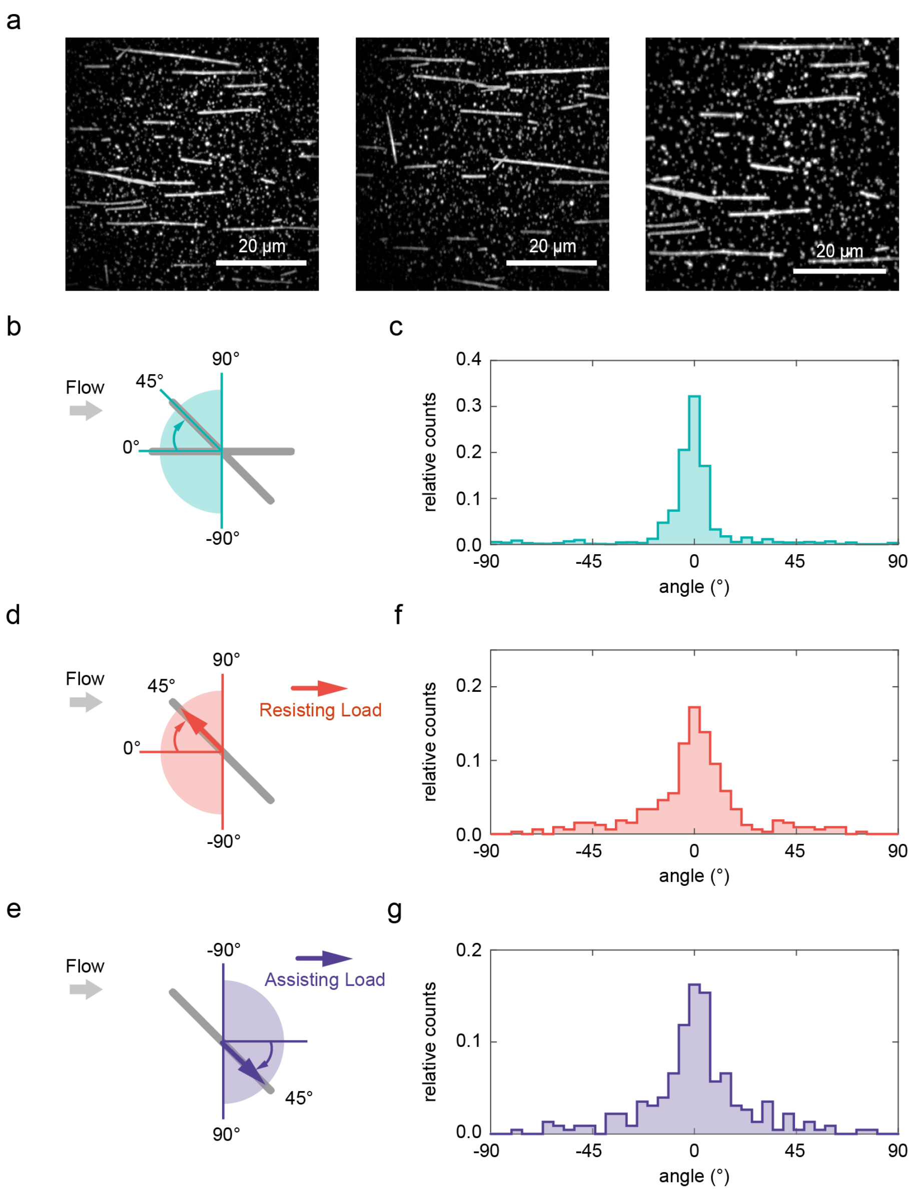


**Figure S3**. The direction of kinesin-1 stepping is dictated by the microtubule orientation. (a) Exemplary fluorescence microscopy images showing the orientation of rhodamine-labeled microtubules in the flow cell. Majority of the microtubules are aligned with the flow direction (from left to right). (b) Schematic representation of the microtubule angular orientation with respect to the flow direction. (c) Length-weighted distribution of microtubule orientations. The angles were assigned based on the fluorescence microscopy images. Data corresponds to one representative experiment in which $n = 427$ microtubules where evaluated. (d,e) Schematic representation of the angle assignment for the motility direction with respect to the flow direction for motors stepping under resisting load (d) and assisting load. (e) Distribution of motor stepping angles for motors stepping under resisting load (f) and assisting load (g) for a representative experiment (number of evaluated molecules $n = 325$ and 228 for f and g, respectively).

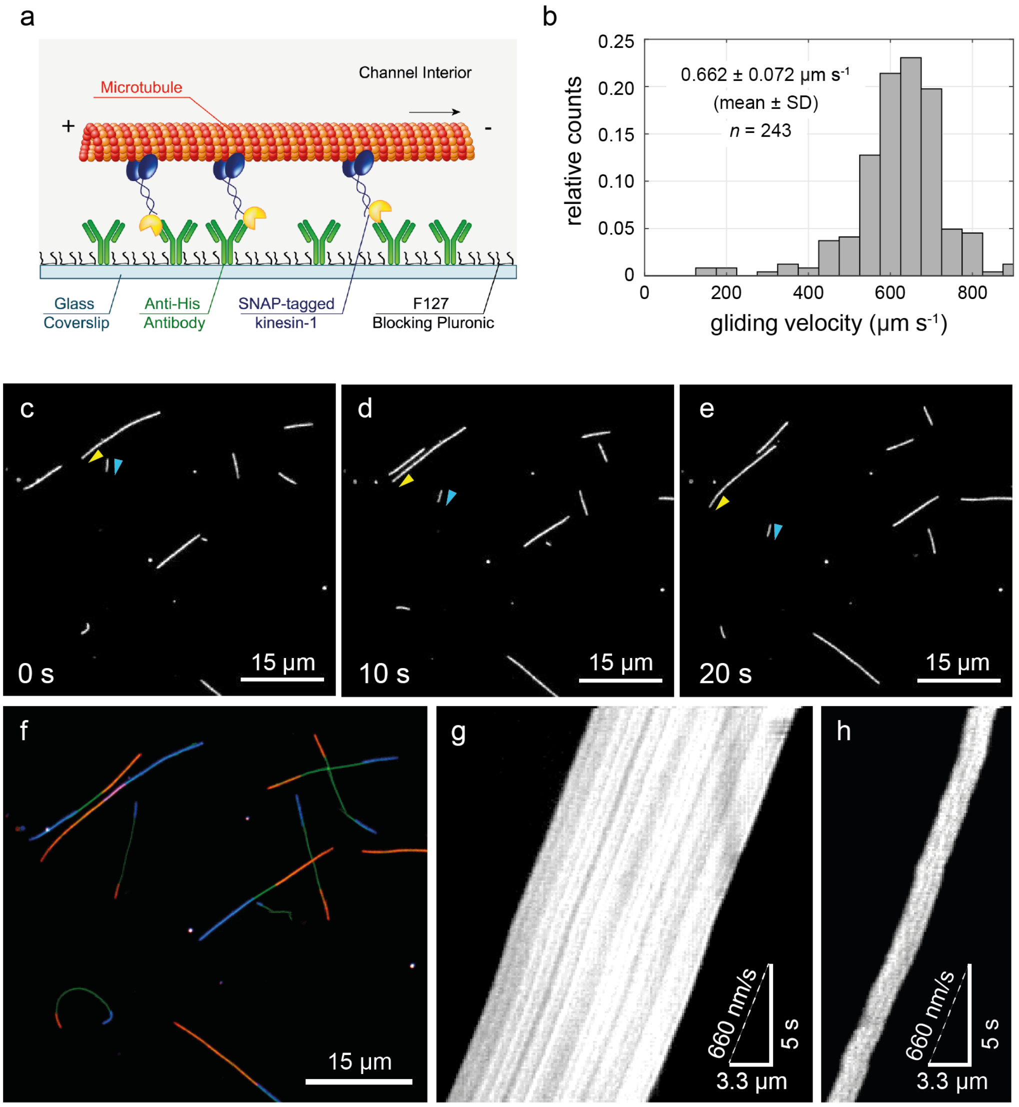


**Figure S4**. Motility of microtubules propelled by SNAP-tagged kinesin-1 motors. a) Schematic overview of gliding motility assay. The microtubule is propelled via surface-bound kinesin motors. The direction of microtubule movement is indicated by the arrow. b) Histogram of gliding velocities based on 243 tracked microtubules. c-e) Fluorescent images of motile microtubules in the sample field of view at times 0 s (c), 10 s (d) and 20 s (e), the arrowheads indicate the direction of gliding for two selected microtubules. f) Three-color overlay of microtubules positions at time 0 s (blue), time 20 s (red) and maximum projection showing the path in between (green). g, h) Kymographs representing microtubule displacement over time. The first kymograph (g) presents the movement of the microtubule indicated by the yellow arrowhead in the panels c-e. The second kymograph (h) represents the movement of the microtubule indicated by the light blue arrowhead.

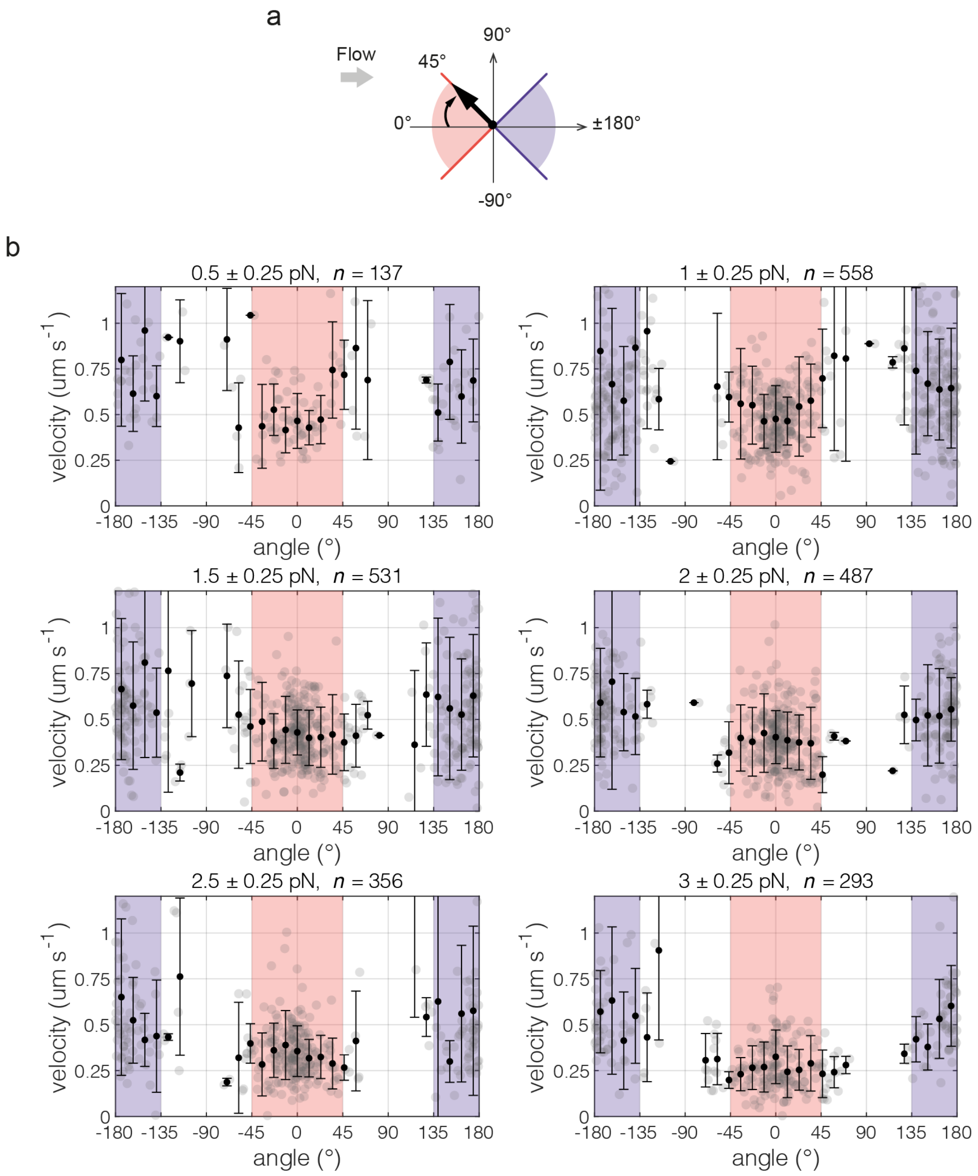


**Figure S5**. Stepping velocities of kinesin-1 as function of stepping angle with respect to the flow direction at different force magnitudes. a) Schematic representation of stepping angle with respect to the flow direction. b) Plots of velocity vs stepping angle for different force magnitudes (indicated above each plot). Events from all experiments were pulled and grouped by force. Black points correspond to mean velocity within a bin of angle values, error bars span standard deviation, gray scatter shows individual events. Red-shaded areas mark angle regions for which motors were considered to step under resisting load, violet-shaded regions mark angle regions for events considered as stepping under assisting load.

a

```
        10         20         30         40         50         60
MADPAECSIK VMCRFRPLNE AEILRGDKFI PKFKGEETVV IGQGKPYVFD RVLPPNTTQE
        70         80         90        100        110        120
QVYNACAKQI VKDVLEGYNG TIFAYGQTSS GKTHTMEGKL HDPQLMGIIP RIAHDIFDHI
       130        140        150        160        170        180
YSMDENLEFH IKVSYFEIYL DKIRDLLDVS KTNLAVHEDK NRVPYVKGCT ERFVSSPEEV
       190        200        210        220        230        240
MDVIDEGKAN RHVAVTNMNE HSSRSHSIFL INIKQENVET EKKLSGKLYL VDLAGSEKVS
       250        260        270        280        290        300
KTGAEGAVLD EAKNINKSLS ALGNVISALA EGTKTHVPYR DSKMTRILQD SLGGNCRTTI
       310        320        330        340        350        360
VICCSPSVFN EAETKSTLMF GQRAKTIKNT VSVNLELTAE EWKKKYEKEK EKNKALKSVI
       370        380        390        400        410        420
QHLEVELNRW RNGEAVPEDE QISAKDQKNL EPCDNTPIID NITPVVDGIS AEKEKYDEEI
       430        440        450        460        470        480
TSLYRQLDDK GTMDKDCEMK RTTLDSPLGK LELSGCEQGL HEIKLLGKGT SAADAVEVPA
       490        500        510        520        530        540
PAAVLGGPEP LMQATAWLNA YFHQPEAIEE FPVPALHHPV FQQESFTRQV LWKLLKVVKF
       550        560        570        580        590        600
GEVISYQQLA ALAGNPAATA AVKTALSGNP VPILIPCHRV VSSSGAVGGY EGGLAVKEWL
       610        620
LAHEGHRLGK PGLGPAHHHH HHHH
```

b

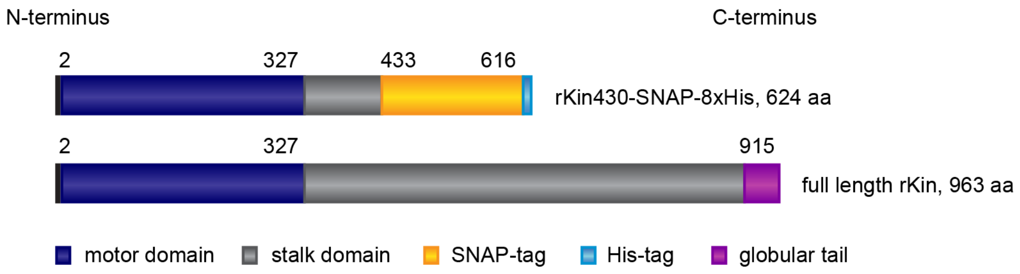


**Figure S6.** The design of kinesin-1-SNAP-tag fusion protein used in this study. a) Full sequence of kinesin-1-SNAP-tag fusion protein (rKin430-SNAP-8xHis) presented in one letter amino acid code. b) Graphical representation of domain localization in kinesin-1-SNAP-tag protein as compared to full length rat kinesin-1 (rKin). The rKin430-SNAP-8xHis protein consists of 624 amino acids. Residues 1 to 430 are residues of truncated rat kinesin-1 (rKin430), where residues 2 to 327 correspond to the motor domain. Residues 433 to 616 correspond to SNAP-tag sequence and the last eight residues, 617 to 624, are histidines used for protein purification.

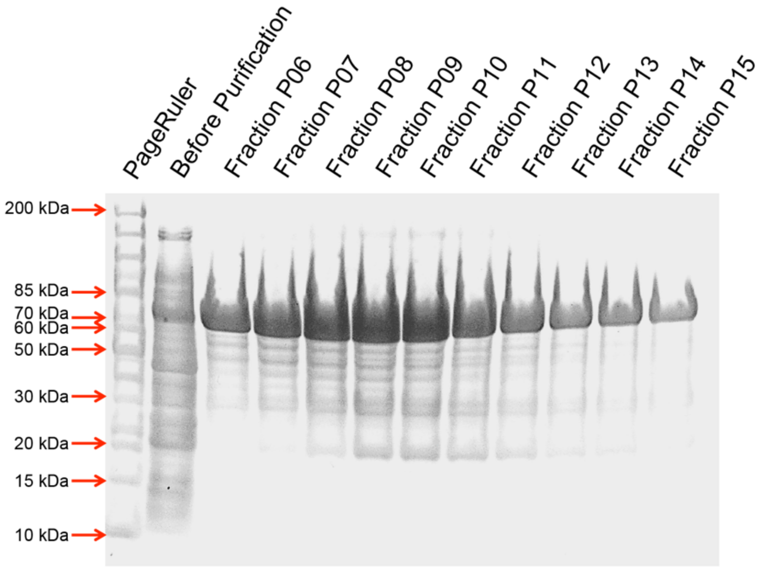


**Figure S7.** SNAP-tagged kinesin-1-rich fractions upon purification resolved in SDS-PAGE. Electrophoresis on 4-12% Tris/Glycine gel (biostep) was carried out for 30 min at 200 V in NuPAGE buffer (Life Technologies). The gel was stained in SimplyBlue Safe Stain solution (Life Technologies) for 20 min and destained in water. For reference, PageRuler Unstained Protein Ladder (Thermo Fisher Scientific Inc.) was loaded in the first lane. The second lane contains the cleared supernatant after cell disruption. The following lanes contain fractions P06-P15 collected upon elution. Fractions P08-P11 were pooled and used for the experiments. The expected size of the expressed protein equals 69 kDa.

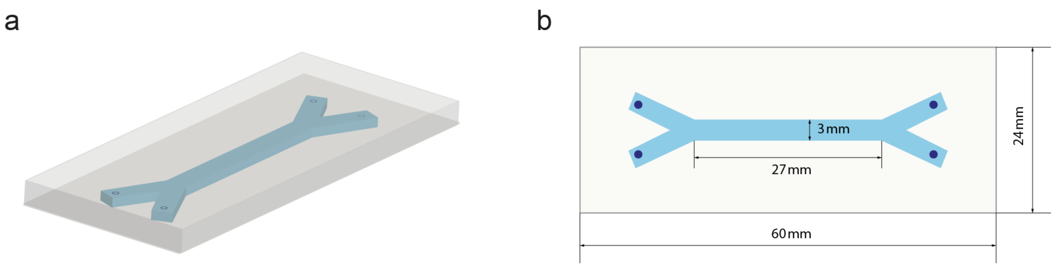


**Figure S8.** Schematics of the microfluidic device used in this study. a) A 3D representation of the microfluidic channel (blue) embedded in a PDMS slab (gray). b) 2D schematics of the chip layout. Dark blue dots indicate holes punched in the PDMS slab for connecting inlet and outlet tubing. One inlet and one outlet were used in a cross configuration during the experiment.

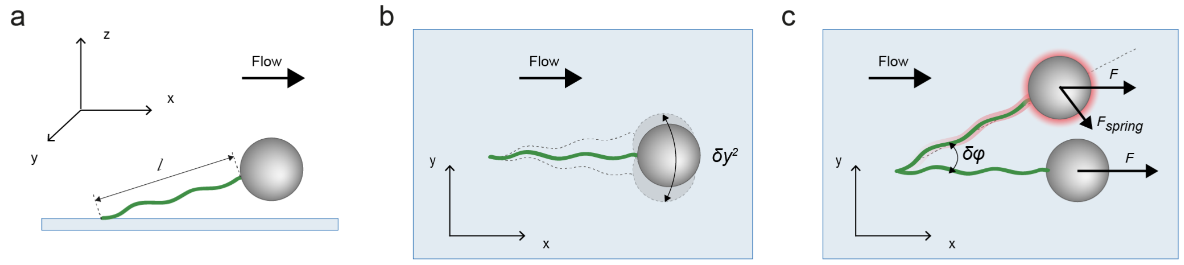


**Figure S9**. Graphical representation of tethered bead fluctuations. a) A bead is tethered to the glass surface via dsDNA of length $l$ (side view - microtubule and motor are omitted for clarity). The direction parallel to the flow direction is defined as x, perpendicular to the flow direction as y and z is defined as the direction out of the glass plane. b) Top view on a fluctuating tethered bead. The displacement in the y direction $\delta y^2$ is indicated by the two-headed arrow. c) Forces acting on the tethered bead in the equilibrium position - the tether is aligned in x direction, and in the out of the equilibrium position - indicated with red shadow (top view). $F$ is the applied drag force, $F_{spring}$ is the restoring force and $\delta\varphi$ is the angle between $F$ and the tether.

**Table S1**. The magnitude of the vertical force component for system geometries used in the microfluidic assay and standard optical tweezers experiments. The vertical force component $F_z$ is related to the force $F_x$ acting in lateral direction in the following manner[1]: $\frac{F_z}{F_x} = \tan\theta \approx \sqrt{\frac{R}{2l}}$, with $R$ – radius of the bead, $l$ – tether length and $\theta$ – the angle between the flow axis and the tether orientation as indicated in **Figure 4**. In the standard optical-trapping assay the bead is attached to the carboxy-terminus of the protein, at the end of the stalk, therefore the length of the tether corresponds to the length of the kinesin stalk and is equal to 60 nm.[2,3]

| | **system characteristics** | | **estimated values** | |
|---|---|---|---|---|
| | bead diameter $2R$ | length of the tether $l$ | inclination angle $\theta$ | vertical force component $F_z$ |
| **microfluidic assay** | 1.08 µm | 16.2 µm | 7° | 13% $F_x$ |
| **optical tweezers** | 0.5 µm[2,3] | 60 nm[2,3] | 55° | 144% $F_x$ |

**Table S2.** Reynolds numbers calculated for conditions used in the microfluidic experiments. Since the width of the channel (3 mm) is significantly bigger than its height (0.1 mm), for the considerations of flow behavior the system can be approximated as two finely spaced infinite parallel plates. For this situation the characteristic dimension of the system ($L$) is defined as twice the distance between the plates;[4] here $L$ = 0.2 mm. $A$ stands for the channel cross section (0.3 $mm^2$). The physical parameters of fluid (density $\rho$ and dynamic viscosity $\mu$) are assumed to be equivalent to those of water at 23°C.

| volumetric flow rate $Q$ (µl $min^{-1}$) | mean flow velocity $v = \frac{Q}{A}$ (mm $s^{-1}$) | Reynolds number $Re = \frac{\rho v L}{\mu}$ (–) |
|---|---|---|
| 10 | 0.56 | 0.12 |
| 20 | 1.11 | 0.24 |
| 30 | 1.67 | 0.36 |
| 40 | 2.22 | 0.48 |
| 50 | 2.78 | 0.60 |

### Supplementary Note 1

##### Correcting the friction coefficient $\gamma$ for surface proximity

Since in our experimental conditions the beads are oscillating relatively close to the surface, it is necessary to correct the friction coefficient $\gamma$ for near-wall effects. For the bead motion in the direction parallel to the surface (in the $xy$-plane, see **Figure S9** and **S10** for the definition of directions), this correction is done by multiplying the Stoke's friction coefficient $\gamma$ by a correction factor $\lambda_{||}$ dependent on the distance of the bead to the surface $z$:

$$\gamma_{||} = \lambda_{||}(z)\,\gamma. \tag{S1}$$

Correction factor $\lambda_{||}(z)$ is defined by Faxen's law[5]:

$$\lambda_{||}(z) = \left[1 - \frac{9}{16}\frac{R}{z} + \frac{1}{8}\left(\frac{R}{z}\right)^3 - \frac{45}{256}\left(\frac{R}{z}\right)^4 - \frac{1}{16}\left(\frac{R}{z}\right)^5\right]^{-1}, \tag{S2}$$

where $R$ refers to the bead radius.

The correction factors $\lambda_{||}(z)$ were determined individually for each flow rate. The distance of the bead from the surface, $z$, was first estimated using steady state force balance (**Figure S10**):

$$\vec{F}_{mag} + \vec{F}_{flow} + \vec{F}_{tether} + \vec{F}_{Br} = 0\,, \tag{S3}$$

where $\vec{F}_{flow}$ is the drag force applied by the flow, $\vec{F}_{mag}$ the force applied by the magnet, $F_{tether}$ is the entropic restoring force of the DNA tether determined by the Equation (13), and $\vec{F}_{Br}$ is the random Brownian force and due to its isotopic nature $< F_{Br} >= 0$.

The equilibrium of forces in $x$ direction can be written as:

$$F_{flow} - cos\theta\, F_{tether} = 0, \tag{S4}$$

where $\theta$ is the angle between the tether and the surface (see **Figure S10**).

The equilibrium of forces in $z$ direction is defined as:

$$F_{mag} - sin\theta\, F_{tether} = 0. \qquad \text{(S5)}$$

From the geometry of the system it follows that $sin\theta = \frac{z}{l+R}$, with $l$ being the extension of the DNA tether. $cos\theta$ can be calculated from the Pythagorean identity $cos\theta = \sqrt{1 - sin^2\theta} = \sqrt{1 - \left(\frac{z}{l+R}\right)^2}$.

Equations (S4) and (S5) can be solved to estimate the $z$ position of the bead and $l$.

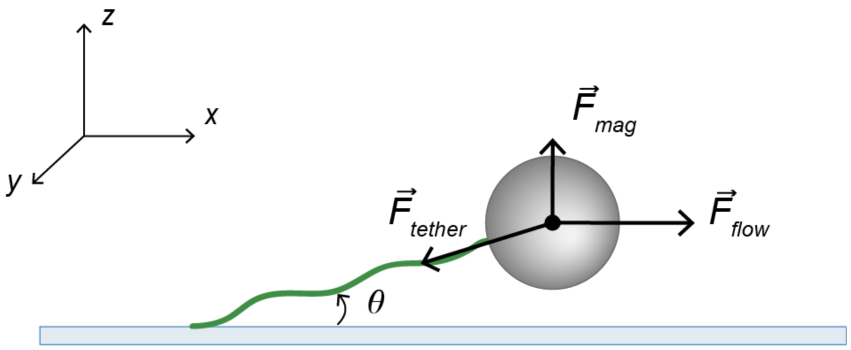


**Figure S10**. Schematics of all forces acting on the bead in the steady state in our assay.

$F_{flow}$ can be estimated using the Stoke's equation, corrected for surface proximity using factor $\lambda_{||}(z)$:

$$F_{flow}(z) = \lambda_{||}(z)\, 6\pi\eta R v(z), \qquad \text{(S6)}$$

where $\eta$ corresponds to the dynamic viscosity of the medium and equals 0.9321 mPa s (for water at 23°C),[6] and $v(z)$ is the height-dependent flow velocity. For the case of laminar flow ($Re < 1$, see **Table S2**) between the parallel plates the flow velocity has a parabolic flow profile described by the following function:

$$v(z) = 2v_{max}\, \frac{z}{h}\left(1 - \frac{z}{h}\right), \qquad \text{(S7)}$$

with the maximum flow velocity $v_{max} = \frac{3Q}{2wh}$, where $Q$ is the volumetric flow rate, $w$ = 3 mm is the width of the flow cell, and $h$ = 100 µm is the height of the flow cell used.

The $z$ positions of the bead and correction factors $\lambda_{||}$ estimated for each flow rate using the force balance and Equation (S2), respectively, are listed in **Table S3**.

**Table S3.** Estimated values of $z$ position of the bead and correction factor $\lambda_{||}$ for each flow rate.

| $Q$ (µl min$^{-1}$) | $z$ position (µm) | $\lambda_{\|\|}$ (–) |
|---|---|---|
| 10 | 1.94 | 1.18 |
| 20 | 1.34 | 1.29 |
| 30 | 1.06 | 1.40 |
| 40 | 0.88 | 1.52 |
| 50 | 0.75 | 1.72 |
